# Liberibacter solanacearum interacts with host psyllid vitellogenin with its membrane proteins

**DOI:** 10.1101/2021.10.29.466487

**Authors:** Poulami Sarkar, Murad Ghanim

**Affiliations:** Department of Entomology, Agricultural Research Organization, Volcani Institute, Rishon LeZion, Israel

## Abstract

*Candidatus* Liberibacter solanacearum (CLso) haplotype D, transmitted by the carrot psyllid *Bactericera trigonica* is a major constraint in carrot production in Israel. Understanding the molecular interactions between the psyllid vector and CLso can facilitate non-chemical approaches for controlling CLso caused-diseases. In this study, we used CLso outer membrane protein (OmpA) and flagellin as baits to screen for psyllid interacting proteins in a yeast-two hybrid assay. We identified psyllid vitellogenin (Vg) protein to interact with both OmpA and flagellin of CLso. As Vg is often involved in innate immunity with its expression tightly linked to autophagy, a major component of the immune response in the cell, we also analyzed the expression of autophagy-related genes to further elucidate this interaction. We used the juvenile hormone (JH-III) to induce the expression of Vg, thapsigargin for suppressing autophagy, and rapamycin for inducing autophagy. The results revealed that Vg negatively regulates autophagy and vice versa. JH-III induced Vg expression significantly suppressed autophagy and, the levels of CLso significantly increased resulting in a significant mortality of the insect. Although the specific role of Vg remains obscure, the findings presented here identify Vg as an important component in the insect immune responses against CLso and may help in understanding the initial molecular response in the vector against Liberibacter.

## Introduction

*Candidatus* Liberibacter solanacearum Haplotype D (CLso) is transmitted by the carrot psyllid *Bactericera trigonica* in Israel (Mawassi et al. 2018; Ghosh et al. 2019; Ghanim et al. 2017). Similar to *Candidatus* Liberibacter asiaticus (CLas), the causative agent of the devastating citrus greening disease (Capoor et al. 1967), CLso is a phloem-limited gram-negative bacterium, transmitted by psyllids in a persistent, propagative manner (Ammar 1994; Ammar et al. 2011; Ghosh et al. 2019; Cooper et al. 2014; Cicero et al. 2017). Many recent studies have addressed the biological and epidemiological relationships between Liberibacter species and psyllid vectors (Sarkar and Ghanim 2020; Ren et al. 2016; Luo et al. 2015; N. Wang et al. 2017; Pelz-Stelinski et al. 2010; Molki et al. 2019), however, little attention has been given to the molecular interactions and the functional validation of Liberibacter and insect proteins that aid in the transmission process. Recently, several studies have unraveled at the transcriptional response of whole psyllids and organs to the acquisition and retention of Liberibacter species (Fisher et al. 2014; Nachappa et al. 2012; Ibanez et al. 2014; Ghosh et al. 2019; Vyas et al. 2015). These studies have shown that the acquisition of different Liberibacter species, by their respective psyllid vectors, induced significant immune responses (Arp et al. 2016; Canale et al. 2016; George et al. 2018; Ammar et al. 2016). Genome and transcriptome sequencing results have recently shown that psyllids do not bear a complete immune response system as has been described in model insects such as Drosophila (Wulff et al. 2014; Lin et al. 2015; J. Wang et al. 2017). Psyllids lack the adaptive immunity and the immune deficiency (Imd) pathway, which generally respond to invasion by gram-negative bacteria, thus leading for example to the ability of Liberibacter species to invade tissues in psyllids where they are able to replicate (Arp et al. 2017; Vyas et al. 2015; Arp et al. 2016). Such findings raise the hypothesis that psyllids use alternative immune responsive mechanisms for combating with the effects of invasion by the bacterium into host cells. On the other hand, psyllids bear an innate defense mechanism against pathogens, which involves both cellular and humoral immune responses (Gill et al. 2017; Arp et al. 2017, 2016). Cellular responses include phagocytosis and humoral responses involve secretion of several antimicrobial peptides (Rosales and Vonnie 2017; Arp et al. 2016). The first line of defense involves recognition of conserved elicitors, molecules or essential structures often known as microbe- or pathogen-associated molecular patterns (MAMPs or PAMPs) by host pattern-recognition receptors (PRR). Bacterial outer membrane proteins (Omp), flagellins and pili appendages are some of the known bacterial virulence factors involved in pathogenesis that elicit immune responses in the host (Vila-Farrés et al. 2017; Liang et al. 2010; Vallet-Gely et al. 2008). OmpA is a major unique integral membrane protein with amphipathic β-barrels which is often involved in cell adhesion and virulence (Bunpa et al. 2020; Weiser and Gotschlich 1991; Qian et al. 2007; McClean 2012). Several human pathogenic bacteria such as *Escherichia coli* (Prasadarao et al. 1996; Maruvada and Kim 2011), *Salmonella enterica* (Singh et al. 2003), *Leptospira interrogans* (Ristow et al. 2007; Zhang et al. 2010), *Neisseria gonorrhoeae* (Makino et al. 1991) also have a direct role of ompA in virulence upon infection in the host cells (Weiss et al. 2008; McClean 2012). The fat body in insects is one of the major immune-responsive organs, where host PRRs against bacterial virulence factors are produced and then directly released into the hemolymph (Feldhaar and Gross 2008). Additionally, hemocytes act as macrophages that have phagocytic activity but also require the presence of PRRs for presenting the pathogen to these macrophages.

One of the major known PRRs is apolipoprotein or vitellogenin (Vg) which belongs to the large lipid transfer protein (LLTP) superfamily having opsonin activity (Babin et al. 1999; Wang et al. 2019; Li et al. 2008). LTTPs consist of a large phosphoglycolipoprotein and a major egg yolk protein precursor (YPP) in insects. They are large molecules (200 kDa) synthesized in the fat bodies and midguts, transported through the hemolymph and sequestered by ovaries with the help of vitellogenin receptors (VgR) via receptor-mediated endocytosis and is subsequently cleaved to generate the nutrient yolk protein vitellin required for the developing oocytes (Zhang et al. 2011; Tufail et al. 2014; Brumin et al. 2020). Although initially, Vg was considered as a female-specific protein, males and sexually immature animals have also been shown to express Vg indicating several roles beyond the nourishment of developing oocytes (Huo et al. 2018; Tufail et al. 2014; Roy-Zokan et al. 2015). It provides host innate immunity with multifaceted functions during several extraneous factors, including chemical exposure, nutritional stress, and infection (Yao et al. 2018; Li et al. 2008). Insect Vg often acts as a pattern recognition molecule to recognize pathogens, enhances macrophage phagocytosis and autophagy, neutralizes viruses by creating cross-links between virions, and often kills bacteria by interacting with the lipopolysaccharides and lipoteichoic acid present in bacterial cell walls (Zhang et al. 2011; Li et al. 2009; Huo et al. 2018; Brumin et al. 2020; Zhang et al. 2005). For instance, silkworm apolipoproteins inhibits *Staphylococcus aureus* by binding to cell surface lipoteichoic acids (Hanada et al. 2011; Omae et al. 2013). Mosquito Vg has been reported to have antiparasitic response against plasmodium (Rono et al. 2010). Bacterial membrane proteins, flagella and pili often serve as PAMPs and immune elicitors that interact with Vg and act PRRs which induce autophagy and trans-generational immune priming (McClean 2012; Confer and Ayalew 2013; Shi et al. 2019; Tong et al. 2010; Salmela et al. 2015; Tetreau et al. 2019; Harwood et al. 2019).

In this study, we show that OmpA and flagellin of CLso interact with Vg of *B. trigonica* and induce autophagy in the psyllid cells. While both Vg and autophagy are important in the immune response against CLso, each seem to negatively regulate the other, and both are important for regulating CLso titers, besides oocyte development, oviposition and egg viability. The described immune responses in this study are crucial for CLso persistence in the insect and seem to be part of a larger mechanism regulated by both the insect and CLso for maintaining the balance between the vector and the pathogen.

## MATERIALS AND METHODS

### Maintenance of psyllid and Liberibacter

CLso infected (CLso+) and CLso free (CLso-) psyllids were maintained on 2 months old Parsley (*Petroselinum crispum*) in separate rooms, under 14 h photoperiodic light at 25 ± 2°C.

### Plasmid vectors

For protein expression studies, we used pRSET-A and pFN2A Flexi vectors (Thermo Scientific) with competent BL21 (DE3) and DH5α cells (NEB, USA). pGAD-T7 Rec (Clontech) was used for the cDNA library preparation, pGADT7-AD (Clontech) as prey vector for one to one assays and were transformed in Y2H Gold yeast cells. pGBKT7 (Clontech) was used as a bait vector (DNA-BD) and was transformed into Y187 yeast cells.

### Yeast two hybrid (Y2H) bait constructs

Sequences of Liberibacter Outer membrane protein (OmpA) and Flagellin (Flg) were derived from the full genome sequence of Candidatus *Liberibacter solanacearum* (Haplotype D) with accession PKRU02000006.1 (Katsir et al. 2018). Full-length coding sequence of OmpA and Flg were amplified from Liberibacter infected (CLso+) psyllids using Q5 DNA polymerase (NEB, USA), cloned into the bait vector-pGBKT7 (EcoRI/BamHI) using In-Fusion HD cloning kit (Takara) and screened for positive recombinants in DH5α cells. Recombinant OmpA-pGBKT7 and Flg-pGBKT7 were finally transformed into Y2H Gold yeast strain separately using Yeastmaker yeast transformation system (Takara, Clontech).

### Psyllid library construction

Total RNA was extracted from around 100 CLso+ psyllids using Trizol (Sigma) and purified using RNAeasy kit (Qiagen). First strand cDNA was synthesized with 3.6 μg of total RNA using Make your own ‘mate & plate’ library system (Takara, Clontech) according to the manufacturer’s instruction. The first stranded cDNA was next amplified to produce double stranded cDNA in 20 amplification cycles by long distance PCR using the Advantage 2 polymerase mix (Takara, Clontech) following the manufacturer’s instructions. The double stranded cDNA was purified using Chroma Spin+TE-400 to eliminate any products below 200bp. The purified ds CDNA was finally co-transformed with pGAD-T7 Rec into competent Y187 using Yeastmaker yeast transformation system (Takara, Clontech) and plated on SD-Leucine agar media. The plates were incubated at 30°C for 3-5 days. Around 2.6 million independent cDNA clones were obtained and the colonies were pooled using YPDA freezing media and stored in aliquots in -80°C.

### Y2H assays

The two baits, OmpA-pGBKT7 and Flg-pGBKT7 were screened against the psyllid library individually, following Matchmaker Gold yeast two-hybrid user manual (Takara, Clonetech). A culture of the bait (Y2HGold) was allowed to mate with psyllid library for 24 h and after mating, the cells were plated on QDO (SD-ATLH) media and incubated at 30 °C for 8-10 days.Developed colonies were re-streaked onto QDO/X-gal^+^ plates to screen for the development of blue color for the β–galactosidase activity, and finally the blue colonies were further streaked onto QDO/Xgal/Aureobasidin (40μg/ml). Plasmids were isolated from the colonies as previously described (Hoffman and Winston 1987) and sequenced for identity.

### RNA extraction, qRT-PCR analysis and Liberibacter abundance

Single psyllids/ guts /ovaries were used for RNA and DNA extraction, both from the same sample using CTAB (Doyle et al. 1987) and as previously described (Sarkar et al. 2020). The guts and ovaries were washed thrice with PBS to remove any contaminants from the hemolymph before proceeding with RNA/ DNA extraction. Final eluted nucleic acid was divided into two aliquots, one for RNA and one for DNA. DNA contaminations from the total RNA were removed with DNaseI (Thermo) and used for cDNA synthesis using Verso cDNA synthesis kit (Thermo) following manufacturer’s instructions. RNA was removed from the DNA sample using RNaseI (Thermo) and was used to measure relative Liberibacter titre using qPCR. Real Time analyses were carried out using 2x Absolute Blue SYBR mix (Thermo) and 1 μl of diluted cDNA in a final volume of 20 μl. Threshold Ct values were calculated in StepOne real Time PCR system (Applied Biosystems) and normalized using the house keeping genes (elongation factor). PCR efficiencies of all new primers were tested using a standard curve, and differential gene expression were analyzed using 2^−ΔΔCT^ quantitation methods (Livak and Schmittgen 2001). The primers used in this study are listed in Table1. Statistical analyses of all qRT-PCR data were conducted by one-way ANOVA with Tukey’s post hoc test (p<0.05). The accumulation of CLso in the hemolymph was also quantified with the method previously described (Sarkar et al. 2020). Statistical analysis was done by Student’s t-test (p<0.05).

### Recombinant protein expression, in vitro translation and pull down assay

#### Prey (Vitellogenin)

Full-length coding sequence for Vg-VWD domain was cloned into pRSET-A, and finally transformed into competent *E. coli* BL21 (DE3) cells to express 6xHis-Vg-VWD. The transformed clones were grown overnight in liquid LB at 37 °C with agitation (200 rpm). A fresh media of 5 ml was seeded with 200 μl of this culture and grown for 4 h or until it reaches 0.6 optical density. At this point, a final concentration of 1 mM IPTG was added and was incubated with agitation at 30 °C for additional 5 h. The cells were finally harvested at 6000 rpm followed by protein extraction. The cells were re-suspended in 200 μl of B-PER along with 1μg of Lysozyme (Sigma-Aldrich) and DNase (Thermo), incubated for 15 min with shaking at room temperature and centrifuged at a high speed for 10 min. The supernatant was collected as purified protein and was used for further protein assays.

#### In vitro translation of Bait protein

Full length CDS of OmpA/Flg was cloned into pFN2A (GST) Flexi vector separately for *in vitro* translation. Briefly, Liberibacter OmpA was amplified using SgfI and PmeI restriction sites (Table 1) and was digested using Flexi Enzyme Blend (Promega). Similarly, pFN2A was also digested and finally ligated to OmpA/Flg using T4 ligase (Thermo). The ligation mixture was used to transform E. coli DH5α for screening a positive recombinant pFN2A-OmpA/ pFN2A-Flg vector. Following this, the recombinant plasmid was isolated and sequenced for confirmation. For *in vitro* translation, we used TNT Quick coupled transcription/Translation System (Promega) following the manufacturer’s instructions.

**Table 1.**
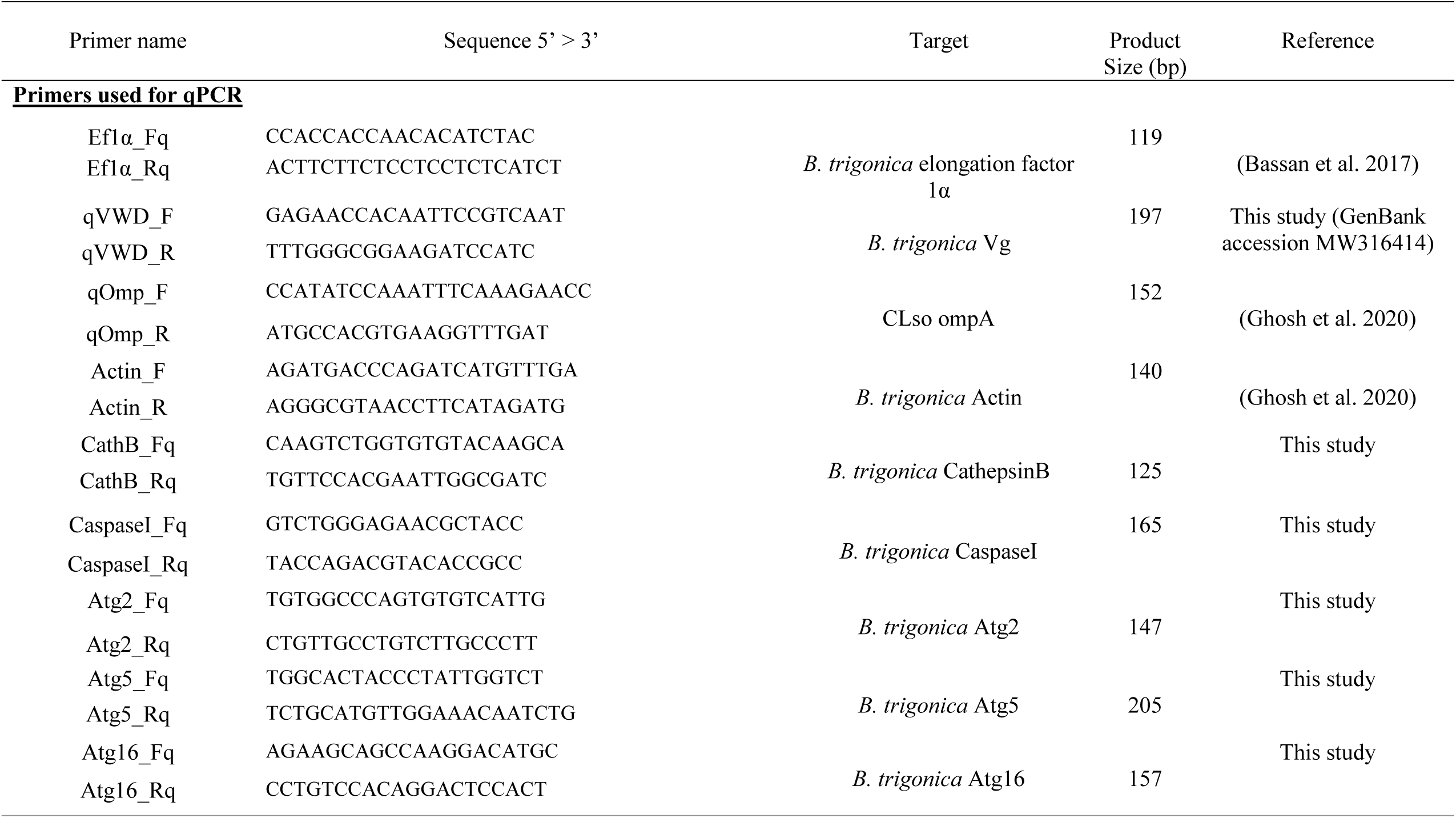
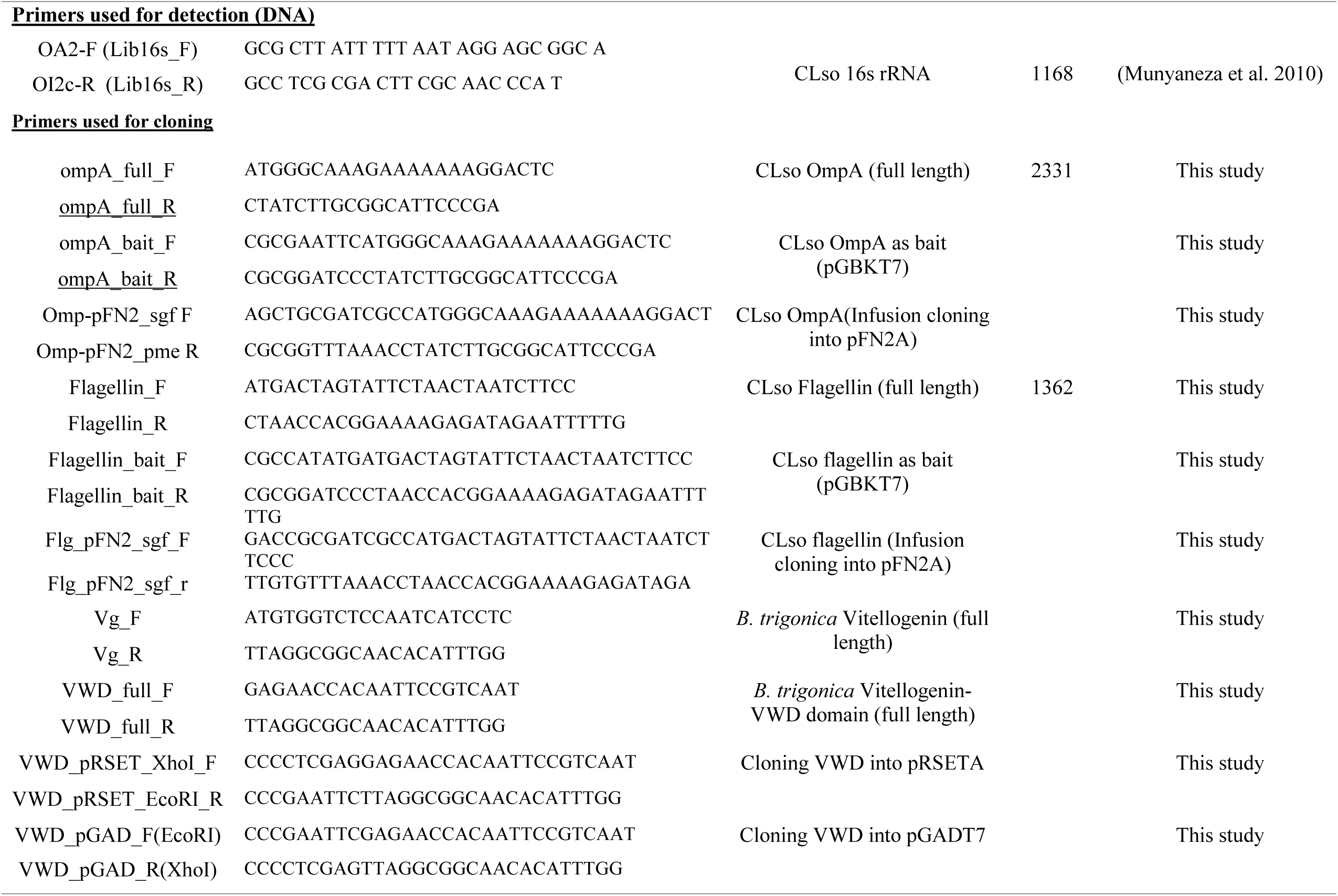
Primers used for PCR detection, qRT-PCR and dsRNA synthesis.

#### Pull down and Western Blot assay

The vector pFN2A was genetically modified to remove the Barnase gene and to express just the GST as control. For pull down assays, we used Magne-GST Pull-Down Systems (Promega, USA). The bait, pFN2A-OmpA/pFN2A-Flg and control GST was immobilized onto Magne-GST particles following the manufacturer’s instructions. Total soluble fractions from prey protein lysate (GST-Vg-VWD) was incubated with the bait immobilized to Magne-GST particles for capture. After washing, the bound proteins were finally eluted by boiling in 1x SDS buffer, separated on 10% SDS PAGE for analysis and detected by western blot using monoclonal anti-polyHis antibody (Sigma Aldrich, Israel).

#### Homology modelling and in silico studies

The open reading frames of the sequences derived from the Y2H assays were annotated using BLASTp and NCBI Conserved Domain Database search (Marchler-Bauer et al. 2015) databases and checked for in-frame reading sequences. Structural analysis for domain identification for was done by Pfam (El-Gebali et al. 2019) and NCBI-CDD. Full-length VWD domain was amplified from carrot psyllids and was used for all further studies. For phylogenetic studies, CLso Vg sequence was aligned with 20 other insect Vgs in Mega 7.0 software with arachnid Vg as an outgroup. The phylogenetic relationship was assessed in CIPRES gateway using Mr.Bayes XSEDE tool with fixed LG+G substitution model and 1 million generation. The tree was finally edited in Figtree program v1.4.4 (http://tree.bio.ed.ac.uk/software/figtree). 3D model structure for Vg was generated by iTasser (Yang and Zhang 2015) with the highest C-score. All known full-length sequences of OmpA/Flg from Liberibacter species were aligned in Mega7.0 (Kumar et al. 2016) and similarity scores with consensus sequences were obtained in ESPript 3.0 (Robert and Gouet 2014). 3D model structure for OmpA was generated by Swiss-Model Tool (Waterhouse et al. 2018) and the server Orientations of Proteins in Membranes (OPM) (https://opm.phar.umich.edu/) and for Flg by Swiss-Model.

### Juvenile hormone (JH-III) hormone treatment

JH-III (Sigma, Israel), which is the principle regulator of Vg synthesis in Hemipterans, was dissolved in ethanol at a concentration of 5μg/μl. To induce the expression of vitellogenin, JH-III was applied to a flush of parsley in an incubation box with 20 female psyllids for 16h. Ethanol was used in the control set of experiment. The psyllids were collected and were used for oviposition, DNA/RNA isolation and immunostaining analyses. The experiments were done in triplicate with minimum of six samples each for qRT-PCR/q-PCR.

### Induction and repression of Autophagy

Autophagy was induced by treating the psyllids with Rapamycin (Sigma, Israel) (a potent mTORC1 inhibitor). The experiment was set up similar to the dsRNA experiments as mentioned previously (Sarkar et al. 2020). Fresh leaf flush was placed in a micro-centrifuge tube, applied with 10 μM of Rapamycin (dissolved in ethanol). 20 female psyllids were released into each jar containing the leaf flush and were allowed to feed for 24 h. Similarly, Thapsigargin (Enco, Israel) was used for autophagy inhibition at a concentration of 10 μM and the application was similar to that of JH-III. Ethanol was used as control. DNA/RNA was extracted from the psyllids, midguts and ovaries for qPCR and qRT-PCR analyses. Midguts were also used for immunostaining. Each experiment was conducted thrice with minimum of six samples each time with a total of minimum 18 samples.

### Immunolocalization

Immunostaining for Vitellogenin, CLso, and auto-lysosomes were done according to the protocol described previously (Sarkar et al. 2020). Psyllid midguts/ovaries were dissected out in PBS, fixed in 4% paraformaldehyde, treated with TritonX-100 and incubated in 1.5% blocking buffer for 1h. Following this, the guts/ ovaries were incubated with Anti-Vg antibody (Abcam) or Anti-OmpB antibody for 1.5 h followed by secondary antibody conjugated with Cy3/Cy5 counterstained with DAPI. The colocalization of CLso and Vg was validated using the Colocalization Finder plugin of ImageJ with Pearson’s correlation coefficient (R value). LysoTracker Green DND-26 (Invitrogen) was used to locate auto-lysosomes according to the manufacturer’s instructions. At least eight midguts were used for each immunolocalization experiments to confirm the consistency of the results obtained. The differences in the signals for auto-lysosomes in CLso- and CLso+ midguts were validated using ImageJ software.

### Oviposition

After JH-III treatment, the number of laid eggs was counted on each leaf flush and were monitored for hatching. Eggs from CLso+ psyllids treated with ethanol were used as a control. To test the presence of transovarial transmission, the leaf flush (for control and JH-treatment) along with oviposited eggs were washed for 2min in 0.5% bleach followed by 50% alcohol and finally washing in sterile distilled water thrice. The eggs were then carefully separated from the leaves from the pedicel with a sterile blade under a magnifying glass and placed on sterile CLso uninfected leaves on a Petri dish (Fig. S1) and was incubated in the plant growth room. The newly hatched nymphs were allowed to feed on the uninfected leaves and were tested for CLso in the 3^rd^ instar stage. For control experiments, the eggs were placed on sterile CLso infected leaves. The differences were analysed by Student’s t-test (p<0.05).

## RESULTS

### Liberibacter interacts with host vitellogenin VWD domain

Liberibacter OmpA contains a surface antigen domain, while Flg acts as a virulence factor. Both were used to screen for interacting proteins in the psyllid vector. A cDNA expression library was prepared using whole psyllids and was mated with the full length CDS of OmpA/Flg expressed as a fusion protein with GAL4-DNA binding domain in Y2HGold that binds to promoters of four reporter genes (AUR1C, HIS3, ADE2 and MEL1). Around 54 isolated colonies were observed in QDO plates for OmpA and 62 for Flg which were re-streaked on QDO/Xgal and QDO/X-gal/Aba plates for confirmation of β-galactosidase activity. Four out of all the colonies for OmpA and three for Flg were identified as parts of VWD domain of Vitellogenin (Vg) using DNA sequencing. To verify the Y2H interaction, full Vg-VWD domain was amplified separately from psyllid DNA, cloned into pGAD-T7 vector and was screened against OmpA/Flg once again which also showed strong β-galactosidase activity (Fig. 1A).

**Fig. 1.**
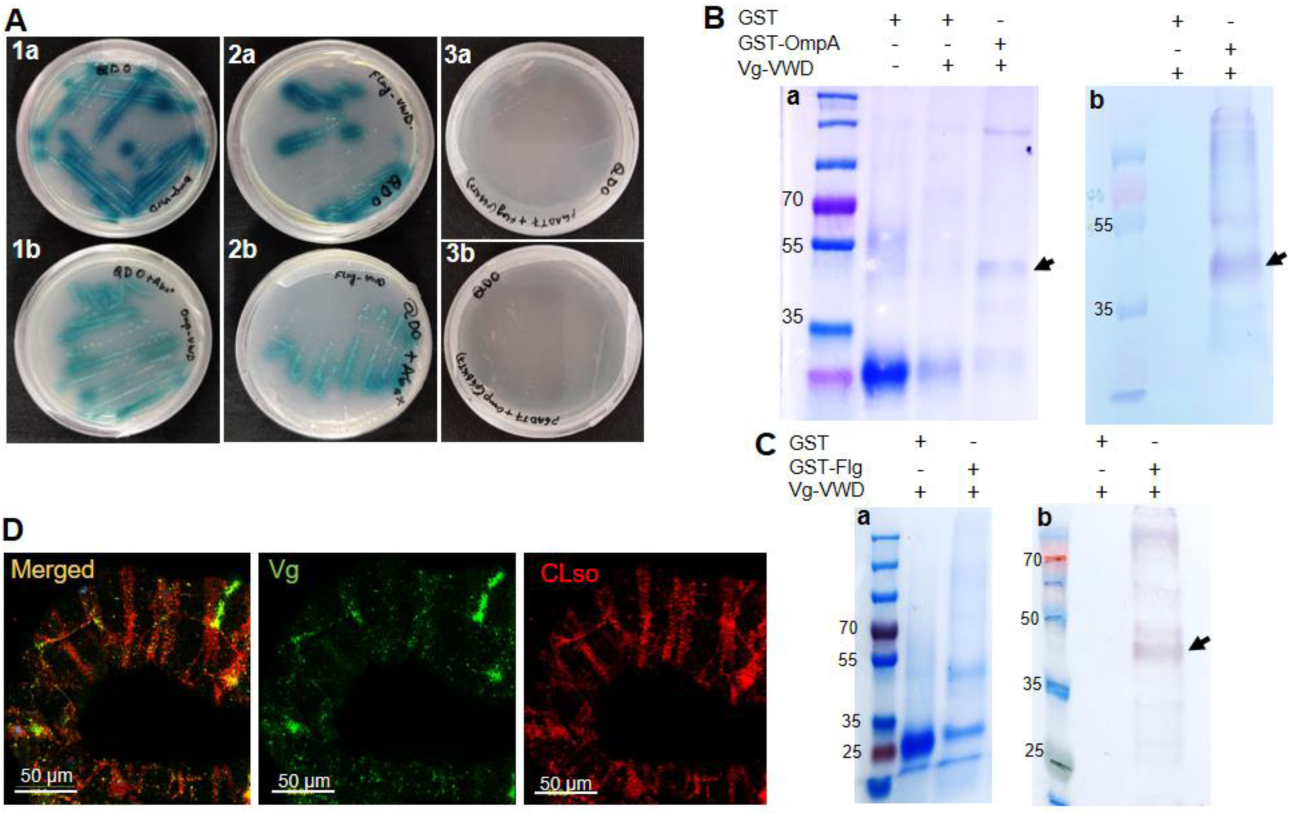
Interaction between Liberibacter membrane proteins and host vitellogenin. **A**, Yeast-two hybrid assay showing strong interaction between Vg-VWD domain and bacterial OmpA (1a, 1b) and Flagellin (2a, 2b) in QDO+Xgal (a) and QDO+Xgal+Aba plates (b). Subsets 3a and 3b show negative control showing no interaction between empty pGADT7 vector with bacterial proteins. **B** and **C**, Detection of Vg-VWD with N-terminal His-tag after pull-down assay using GST-tagged OmpA (B) and Flg (C) as baits by SDS-PAGE (a) and western blot (b) using anti-His antibody. **D**, Immunostaining of CLso+ midguts with anti-Vg antibody (green) and anti-CLso antibody (red) showing spatial co-localization of the two (yellow).

Interaction between Vg and OmpA/Flg was further confirmed using a pull-down assay using OmpA as a bait. A band of approximately 48 kDa was observed in SDS-PAGE as well as western blots using Anti-His antibody when OmpA and Vg were included in the assay (Fig. 1B) or when Flg and Vg were used (Fig. 1C). No band was detected when GST control was used as a bait, indicating a specific interaction between OmpA/Flg and Vg-VWD. To further confirm the interaction, spatial localization of Vg was performed in dissected midguts from CLso infected psyllids using immuno-localization with specific antibodies for Vg and CLso. The signal observed in the midgut indicated a partial overlap in the fluorescent signals of Vg and CLso, which indicated a physical proximity in midgut cells (Fig. 1D).

### In silico analyses

Only one Vg homolog was identified from the psyllid transcriptome (Ghosh et al. 2019). The coding sequence was validated by cloning and sequencing. Structural analysis of Vg revealed three major domains, which include Lipoprotein_N terminal domain (LPD_N), a 1943 domain (DUF1943) of unknown function and C-terminal von Willebrand factor type D domain (VWD) that are usually found in conventional Vgs/LLTPs (Fig. 2A). Phylogenetic analysis showed that the carrot psyllid Vg clustered in the same clade as two other psyllids with 90.7% identity with the potato psyllid *Bactericera cockerelli*. It also clustered in a separate clade formed by other hemipterans in the group.

**Fig. 2.**
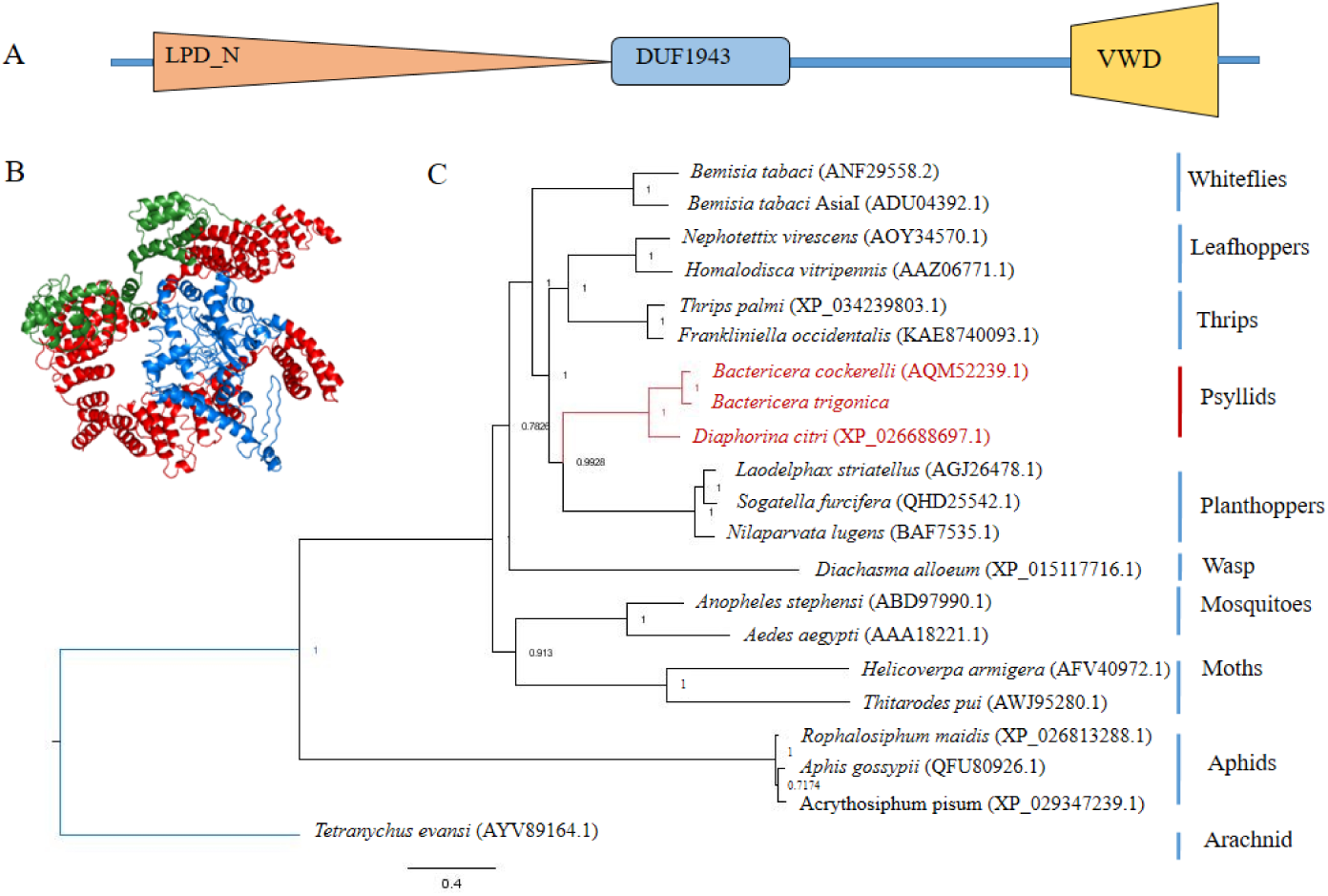
Structure and domain architecture of Vitellogenin. **A**, Vg contains three conserved domains; Lipoprotein LPD_N terminal domain, DUF1943 and VWD domain. **B**, 3D structure of psyllid Vg, modelled by iTasser with highest C-score of 0.32 shows the two major domains; LPD_N (red), and VWD (blue). **C**, Phylogeny of amino acid sequences of all known psyllid Vg proteins showing clustering within Hemipteran clade with *T. evansi* used as an out-group.

Amino acid sequences of all known Liberibacter OmpA and Flg were aligned for sequence identity. Despite an overall high level of identity, conservation of sequences were found to be scattered for both OmpA and Flg. Domain analysis for OmpA revealed four polypeptide transport-associated domain (POTRA) and a bacterial surface antigen. Homology modelling also revealed a 3D structure for OmpA with all the major domains (Fig. S2). Similar analyses for flagellin revealed a signaling domain and a polymerization domain and show minor heterogeneity when aligned with conserved sequences (Fig. S3).

### CLso induces vitellogenin and autophagy-related genes in psyllids

Differential expression profiles of Vg and autophagy related genes were studied in CLso uninfected (CLso-) and infected (CLso+) psyllids. Immunostaining revealed higher expression of Vg in the CLso-infected midguts and its expression was upregulated in CLso+ psyllids as compared with control uninfected ones by 5.9 fold changes in whole body samples and by 1.75 fold changes in the midguts (Fig. 3A and B).

**Fig. 3.**
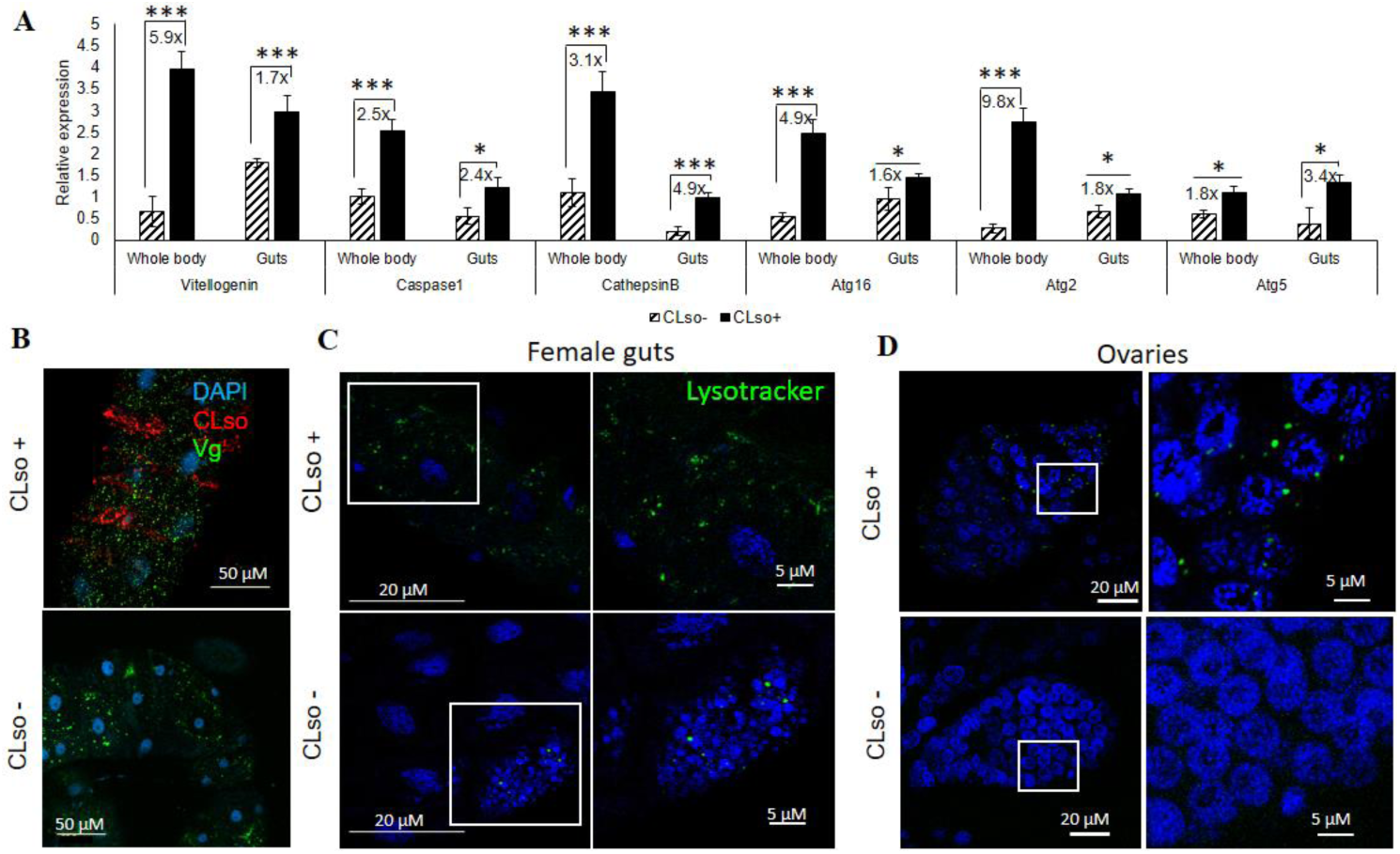
Expression profiles of Vitellogenin (Vg) and the autophagy-related (Atg) genes CaspaseI, Cathepsin B, Atg16, Atg2 and Atg5 in CLso uninfected and infected psyllids. **A**, Relative expression using real-time PCR showing upregulated gene expression of Vg and Atg-genes in CLso+ whole bodies and midguts compared to CLso-psyllids (*,p< 0.05; ***, p<0.001). **B**, Higher expression of Vg (green) in CLso+ midguts compared to CLso-. **C** and **D**, staining of acidic compartments (lysosomes and autolysosomes) using LysoTracker Green showing higher lysosomal activity in CLso+ midguts (C) and in ovaries (D) as compared to CLso-psyllids.

Cathepsin-B and Caspase-I, known immunity genes involved in lysosomal functions against several pathogen invasions, were found to be upregulated in CLso+ psyllid whole body and midguts. The autophagy genes Atg16, Atg2 and Atg5 involved in autophagosome formation were also upregulated in both midguts and whole psyllids (Fig. 3A). Higher lysosomal activity in CLso+ midguts (Fig. 3C) and ovaries (Fig. 3D) as compared to CLso-psyllids was observed when guts cells were stained with Lysotracker, which specifically binds to acidic organelles, indicating higher formation of autolysosomes. The intensity of the Lysotracker signal in CLso+ midguts were 3.12 ±1.2 times more than CLso-as validated using ImageJ software.

### Vitellogenin overexpression impairs the induction of autophagy and vice versa

Vg expression was measured following psyllid treatment with the JH-III hormone, which is the main regulator of Vg production during oogenensis. After 16 h of exposure to JH-III, significant elevation of Vg expression was observed in whole bodies and midguts of both male and female with significantly higher induction in females (Fig. 4A). Vg induced females also had increased number of fat bodies as seen while dissection (data not shown). Induction of Vg also induced Liberibacter titre in the midguts as well as in the hemolymph as measured by qPCR and immunostaining (Fig. 4 B-D), where also elevated autolysosomal activities were observed (Fig. 3). Vg and CLso were seen to mostly colocalize in midguts and ovaries as validated by Pearson’s correlation coefficient (R >0.75). Induction of Vg expression, however, caused a significant downregulation of the autophagy-related genes in whole females (Fig. 5A) midguts (Fig. 5B) and ovaries (Fig. 5C). The presence of autolysosomes was almost negligible in the JH-III treated psyllid midguts (Fig. 5D) and ovaries (Fig. 5E) compared to the control treatments.

**Fig. 4.**
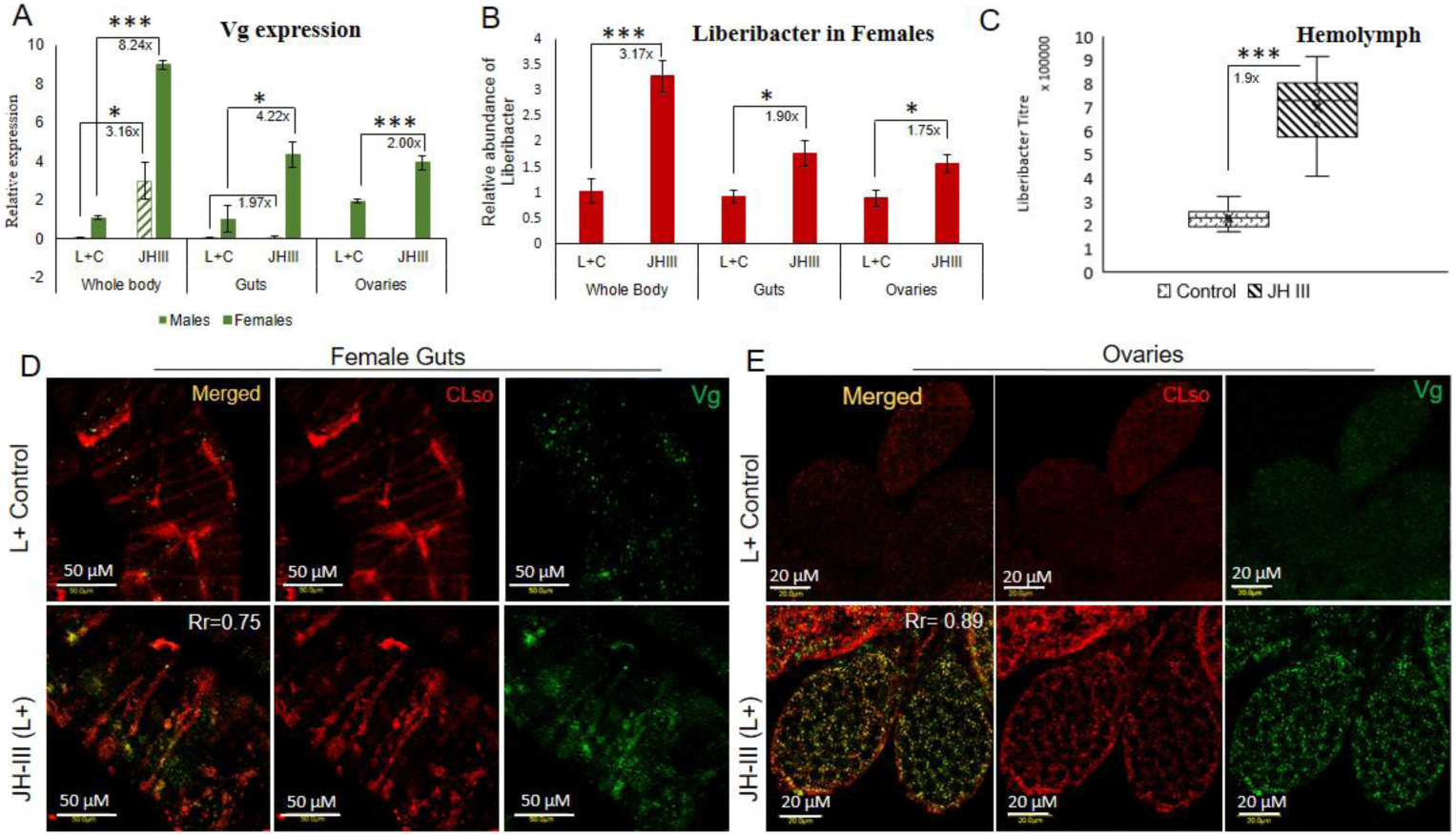
Effect of JH-III hormone on Vg and CLso. **A**, Relative expression of Vg in male and female whole body, midguts and in ovaries showing induced expression of Vg throughout with females showing much higher expression than in males (*, p< 0.05; ***, p<0.001). **B**, Relative titre of CLso (Omp) in female whole bodies, midguts and in ovaries after JH-III application (p≤0.05). **C**, Elevated CLso titre in the hemolymph of JH-III treated psyllids (*denotes p< 0.05; *** denotes p<0.001). The fold change for each gene is mentioned beside the bars. **D** and **E**, Immunostaining of Vg and CLso showing induction of Vg (green) expression and increase in CLso titre (red) upon JH-III application along with their co-localization. The colocalization was validated using ImageJ with Pearson’s correlation coefficient (R value) of 0.75 and 0.89 for guts and ovaries, respectively.

**Fig. 5.**
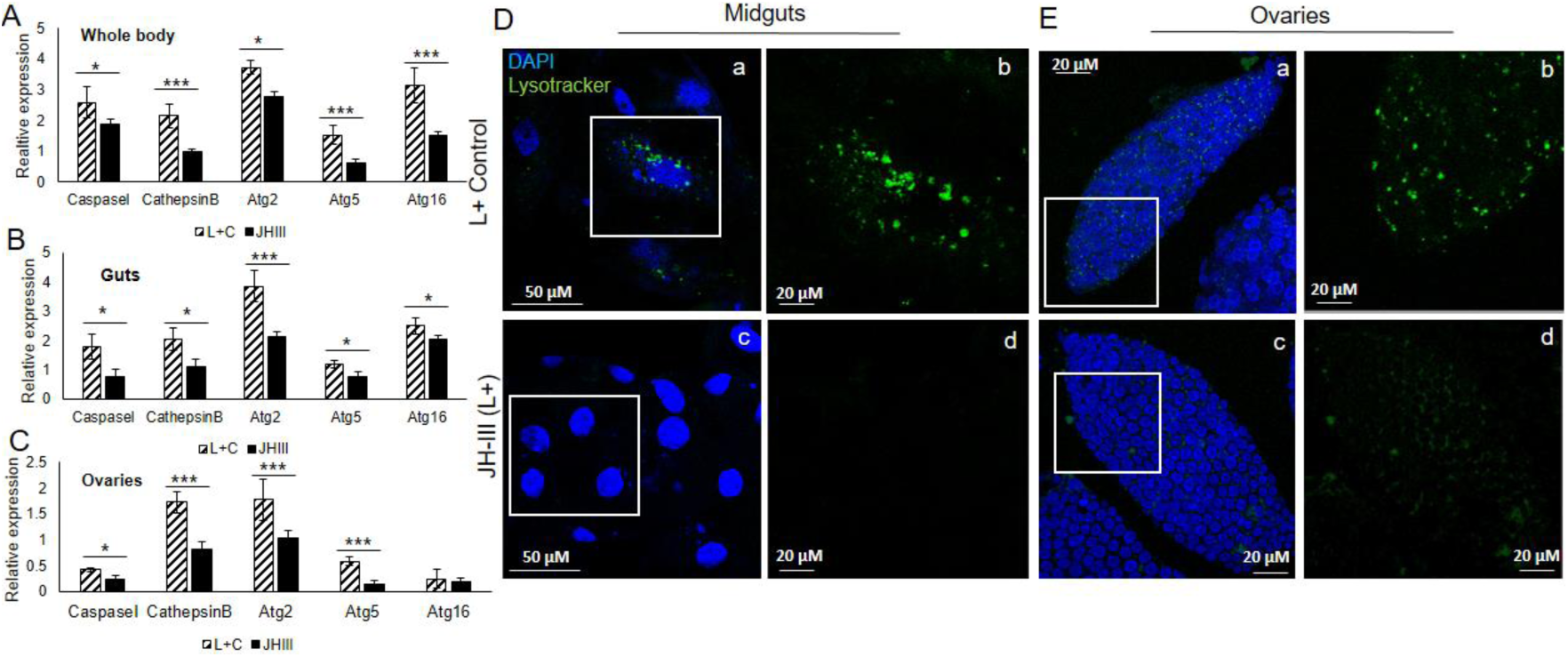
Effect of JH-III on autophagy. Relative expression of lysosomal and autophagy genes in whole bodies (**A**), midguts (**B**), and ovaries (**C**) showing downregulation of all the known genes upon JH-III application (p≤0.05). **D** and **E**, representative images showing reduction of autophagy and lysosomes in the JH-III applied psyllids. Staining of the midguts (**D**) and ovaries (**E**) with DAPI (blue) and lysosomes (green) with b and d showing magnified images of the insets in a and c, respectively. * denote significance p< 0.05 and *** denotes p< 0.01.

Interestingly, application of thapsigargin, that specifically inhibits autophagy, reduced the expression of Vg along with all other autophagy genes, while causing an increase in CLso levels, as seen in qRT-PCR (Fig. 6A) and immunostaining (Fig. 6C). On the other hand, using the specific autophagy inducer rapamycin significantly induced autophagy and autophagy related genes and reduced the expression of Vg and Liberibacter titres in the psyllid midguts (Fig. 6B and 6C).

**Fig. 6.**
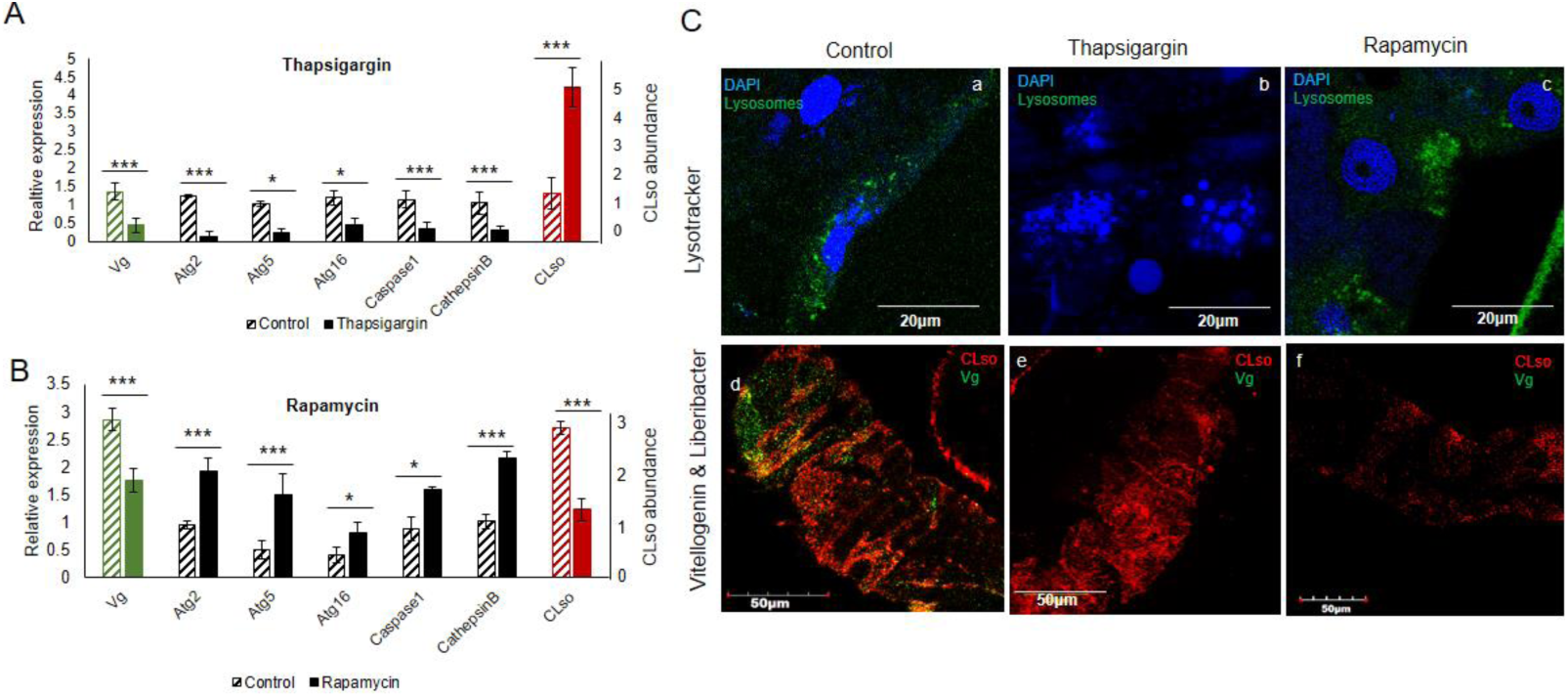
Effect of Thapsigargin and Rapamycin on Vg, CLso and autophagy. **A**, Relative gene expression of Vg, autophagy genes and CLso titre in the psyllid midguts upon Thapsigargin application (p<0.05). **B**, Relative gene expression of Vg, autophagy genes and CLso titre in the psyllid midguts upon Rapamycin application (p<0.05). **C**, Staining of lysosomes (green) and nuclei (blue) showing disintegrated nuclei and absence of autophagy upon Thapsigargin application (b), and increase in lysosomal activity upon Rapamycin application (c) as compared to the control midguts (a). Lower panel showing decrease in vitellogenin and increase in CLso titre upon Thapsigargin application (e) and lower CLso abundance upon Rapamycin application (f) compared to control midguts (a). * denoe significance p< 0.05 and *** denotes p< 0.01.

### Induction of Vg impairs egg development, oviposition and viability

Since Vg has an important role in oogenesis and egg development we investigated its induction following JH-III treatment on mortality, fecundity and fertility, as compared with induction as a result of the presence of CLso. No significant mortality was observed in the JH-III exposed female psyllids as compared to the controls. However, a significant reduction in the number of eggs laid by Vg induced female psyllids was obtained (Fig. 7A). JH-III application further induced oocyte development in female psyllids post 48h treatment with higher number of mature oocytes that was observed as compared to that of control females (Fig. 7B). Moreover, only 3% of the eggs laid by JH-III exposed females hatched compared to 83% viability in CLso+ control eggs (Fig. 7C), with observed malformations and developmental defects in the laid eggs following JH-III treatment (Fig. 7D).

**Fig. 7.**
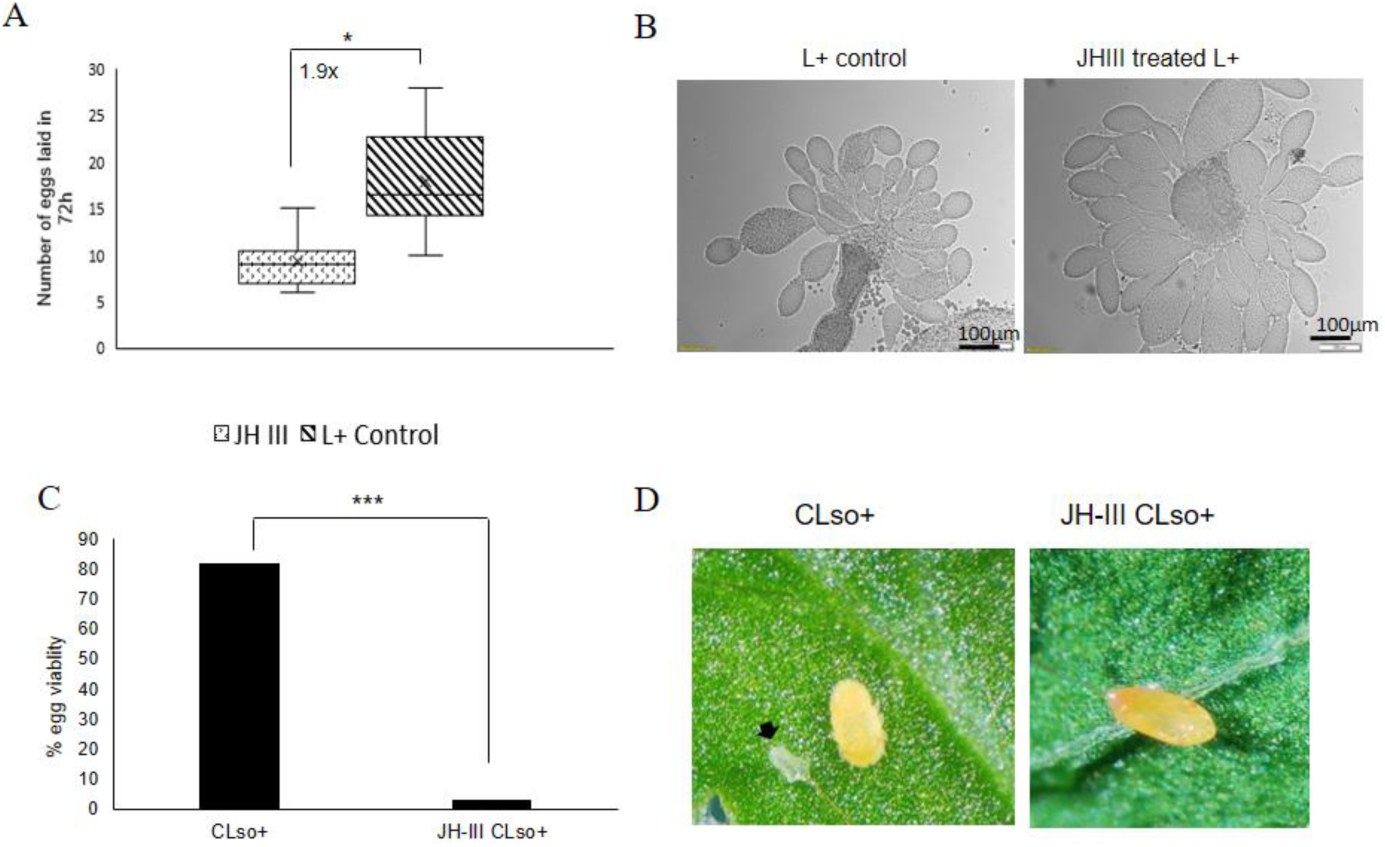
Effect of JH-III on egg development and viability. **A**, JH-III application reduces fecundity (*, p< 0.001). **B**, Representative images showing increased number of ovarioles developing in ovaries dissected from females that were exposed to JH-III. **C**, Number of hatched eggs (fertility) is significantly reduced upon JH-III treatment as compared to control psyllid eggs. **D**, Representative image showing a nymph hatching from CLso+ control egg and a dehydrated egg laid by JH-III treated females. White arrow shows the egg shell from which the nymph hatched.

### Absence of transovarial transmission

Since JH-III significantly induced CLso titers and ovary development in the psyllids, we tested whether these effects may cause the bacterium to be transferred to the developing oocytes by transovarial transmission. One out of 25 hatched nymphs that developed from eggs laid by CLso+ females that were reared on CLso-leaves tested positive for CLso, indicating very low or negligible transovarial transmission. Newly hatched nymphs that developed from eggs laid by CLso+ females that were reared on CLso-leaves, were completely viable and efficiently acquired CLso when fed on CLso+ leaf flush. Interestingly, ovaries dissected from CLso+ females treated with JH-III all tested positive for CLso (Fig. S4), however, laid eggs by CLso+ and JH-III treated females, tested negative for CLso and they were 100% unviable (Fig. 7C and D), suggesting that JH-III treatment indeed accelerates CLso penetrance into ovaries, however this treatment is fatal for the eggs.

## DISCUSSION

Bacterial membrane proteins have virulence properties and often act as pattern recognition molecule which induce host immune response (Liang et al. 2010; Confer and Ayalew 2013). In this study, we used Liberibacter outer membrane protein (OmpA) and flagellin (Flg) as baits to screen for specific insect host proteins that interact with OmpA and Flg. Since both of these bacterial proteins bear an important role in adhesion and virulence, we expected that both proteins will interact with psyllid proteins either for adhesion during the transmission process or for bypassing the host immunity. Yeast-two hybrid assays using OmpA/Flg as baits revealed their interaction with host Vg-VWD domain when screened against psyllid cDNA library (Fig. 1A). Pull-down assays and spatial co-immunolocalization using confocal microscopy also indicated specific interactions between both bacterial proteins and Vg (Fig. 1B-C). Vg has been reported to act as a pattern recognition molecule against pathogens and it has been shown to mediate their degradation by hemocytes through phagocytosis (Salmela and Sundström 2018; Rono et al. 2010; Tong et al. 2010; Singh et al. 2013). Both bacterial proteins also carry immune elicitors and are able to induce immunity in subsequent generation through the process known as transgenerational immune priming (TGIP) (Milutinović and Kurtz 2016; Sheehan et al. 2020; Rosales and Vonnie 2017; Salmela et al. 2015). In this study, only one Vg homolog was identified from the carrot psyllid transcriptome (Ghosh et al. 2019) and the full-length coding sequence was assembled and cloned. It has two major domains, apolipoprotein domain (LPD_N), which helps in lipid transport and VWD domain, a multifunctional domain involved in maintaining homeostasis (Sun et al. 2013; Schneppenheim and Budde 2011). Based on phylogenetic analysis, Vg clustered with the other two psyllid Vgs that clustered within the hemipteran clade (Fig. 2). The surface antigen region of OmpA (Fig. S2) and hypervariable region of flagellin (Fig. S3) are reported to possess adhesion-like properties and they often act as microbe-associated molecular pattern (MAMP) inducing virulence (Ramos et al. 2004; Bunpa et al. 2020; Prasadarao et al. 1996; Qian et al. 2007; McClean 2012).

The known interaction between Vg and bacterial membrane proteins was a trigger to investigate the role of Vg in Liberibacter pathogenesis and host immunity response. The function of vitellogenin is often known to accompany programmed autophagy during development and under stress conditions, and a tight link between the two has been reported in various studies (Li et al. 2009; Seah et al. 2016; Bryant and Raikhel 2011). In this study, we investigated the gene expression profiles of autophagy-related genes (available from the transcriptome) and the presence of autolysosomes in CLso+ psyllid midguts and ovaries as a result of Vg induction. The results showed a significant upregulation of Vg along with the autophagy-related genes (Atg2, Atg5 and Atg16) in CLso+ whole body as well as in midguts of carrot psyllids (Fig. 3A and B). Cathepsin B and caspase I involved in lysosomal activity were also upregulated in CLso+ psyllids (Fig. 3B). The presence of lysosomal bodies and autolysosomes were evidently higher in CLso+ psyllid midguts and ovaries (Fig. 3C and D). Higher number of lysosomes indicate higher lysosomal activity and higher autophagy as autophagosomes deliver cytoplasmic materials or cellular debri to the lysosomes for degradation. These results explains the joint and orchestrated function of both Vg and autophagy-related genes upon Liberibacter infection for maintaining homeostasis, and the crucial role of these functions for maintaining the cell viability. However, when Vg expression was induced with the application of JH-III hormone (Fig. 4), there was a drastic reduction in autophagy and lysosomal activity (Fig. 5), and the expression of autophagy-related genes and lysosomal proteases were significantly downregulated (Fig. 4 and 6).

Moreover, Vg induction drastically reduced oviposition and egg viability (Fig. 7). It is known that overexpression of Vg induces ageing and impairs the induction of autophagy and lysosomal genes required to maintain longevity (Seah et al. 2016) and autophagy is induced during the synthesis phase of vitellogenin (Vg) in the fat body to maintain developmental switches, regulate immunity and recycle cellular components during development (Weng and Shiao 2020; Bryant and Raikhel 2011; Mathieu 2015). On the other hand, upon autophagy arrest by thapsigargin, Vg expression was reduced along with the autophagy-related gene expression in the psyllids (Fig. 6A and C). This result is in congruence with previous reports where it has been shown that cellular calcium plays a regulatory role in Vg production and thapsigargin acts as calcium mobilizing agent while blocking autophagy and inducing ER stress and apoptosis (Yeo and Mugiya 1998; Yeo 1998; Wang et al. 2016; Lindner et al. 2020; Ganley et al. 2011). Induction of apoptosis is also known to activate Perk-eIF2 pathway and Ire-1 dependent decay (RIDD) of mRNA, which results in reduced synthesis and degradation of Vg mRNA, respectively (Luo et al. 2017; Metcalf et al. 2020). Additionally, Liberibacter titers increased significantly and its signal was seen to be diffused in the psyllid midguts treated with thapsigargin. Interestingly, inducing autophagy, by applying rapamycin, reduced Vg expression as well as Liberibacter in the psyllid midguts. The reduction in Liberibacter might be due to increased autophagy. This indicates that both vitellogenesis and autophagy are important for cell survival and are integral parts of developmental process, which help in maintaining cellular homeostasis. Any imbalance between the two may disrupt the homeostasis and may lead to cell death (Seah et al. 2016; Bryant and Raikhel 2011; Raikhel 1986b, 1986a).

The results of this study also show elevated titers of Liberibacter in the JH-III treated midguts as well as in the hemolymph in the absence of autophagy (Figs. 4 and 6). Higher abundance of Liberibacter titer in the midguts and hemolymph suggests a role of Vg in presenting Liberibacter to the cells inducing autophagy, whose absence results in higher titers of the pathogen in the system. There might also be a role for Vg in transgenerational immune priming in CLso+ psyllids and a possibility of transovarial transmission in the absence of autophagy. We could not detect CLso in viable ovaries and laid eggs, which indicate absence of transovarial transmission. Nymphs developing from eggs laid by CLso+ females that hatched and fed on CLso-leaves were also negative for CLso. This implies that nymphs acquire CLso by feeding only on infected leaves with CLso, and not by transovarial transmission. Surprisingly, ovaries dissected from CLso+ females which were treated with JH-III tested positive for Liberibacter in the absence of autophagy. However, induction of Vg reduced egg viability although vitellogenic development in the oocytes and the number of ovarioles was greater as compared to the control CLso+ psyllid ovaries (Fig. 7). This possibly happened due to the lack of autophagy, which disrupted proper cellular development. These results suggest that the ovaries tested positive for CLso as a result of Vg induction or reduction in autophagy. These result suggest a role for Vg in the defense mechanism and might be involved in TGIP as depicted in the model presented in Fig. 8, although an exact mechanism remains unknown.

**Fig. 8.**
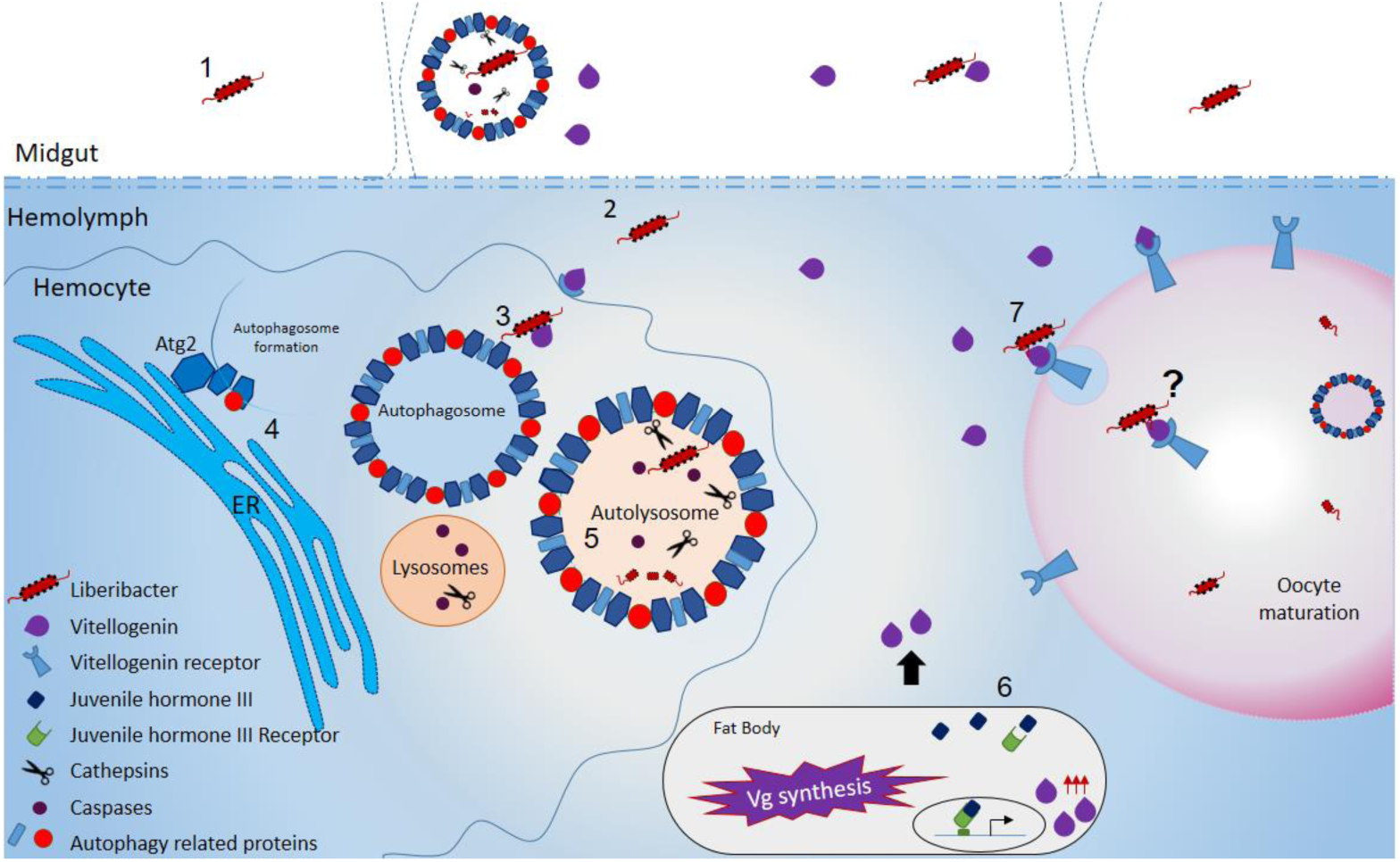
A suggested model depicting how Liberibacter induces vitellogenin production and autophagy with a possible role in transgenerational immune priming. Once Liberibacter is in the midgut lumen (1), it crosses the gut wall cellular barrier by unknown mechanism, possibly receptor-mediated endocytosis, and translocates to the hemolymph (2). In the hemolymph Liberibacter is recognized by pattern recognition molecule vitellogenin (Vg) which in turn induces the host immune response by presenting the pathogen to the autophagosomes (3). The autophagosomes initiated from the ER (4) fuses with lysosomes to form autolysosomes, leading to pathogen destruction with the help of Cathepsins and Caspases (5). Vg, on the other hand, is induced by the Juvenile hormone (JH-III) in the fat bodies and released into the cytosol (7).Upon developmental cues, Vg binds to the Vg-receptor leading to oocyte maturation (7). However, JH-III application under experimental conditions as presented in this manuscript suppresses autophagy, leading to inability of Vg to present Liberibacter to the autophagosomes. This allows Vg-bound Liberibacter to reach and enter the developing oocytes. However, as obtained in control experiments we did not detect transovarial transmission, suggesting that Vg is involved in oocyte development, and possibly in immune priming. The mechanism behind the later however, remains unclear and is denoted by ‘?’.

In summary, the results presented in this study reveal that both vitellogenin and autophagy are essential in regulating Liberibacter transmission in the psyllid and the generated stress responses in the cells following CLso infection, although the exact role of Vg remains unclear. In future studies it will be interesting to investigate whether Liberibacter interacts with vitellogenin to manipulate the host immune response for its survival or it is a host defense mechanism against Liberibacter to reduce cellular stress and maintain homeostasis.

## Acknowledgements

We thank Eduard Belausov for technical help with the confocal microscope and members of the Ghanim laboratory for technical support and for providing comments on preliminary versions of the manuscript text.

This research was supported by grant 1163/18 from the Israel Science Foundation to M.G.

