## Supplementary material for "Liberibacter solanacearum interacts with host psyllid vitellogenin with its membrane proteins": Fig. S

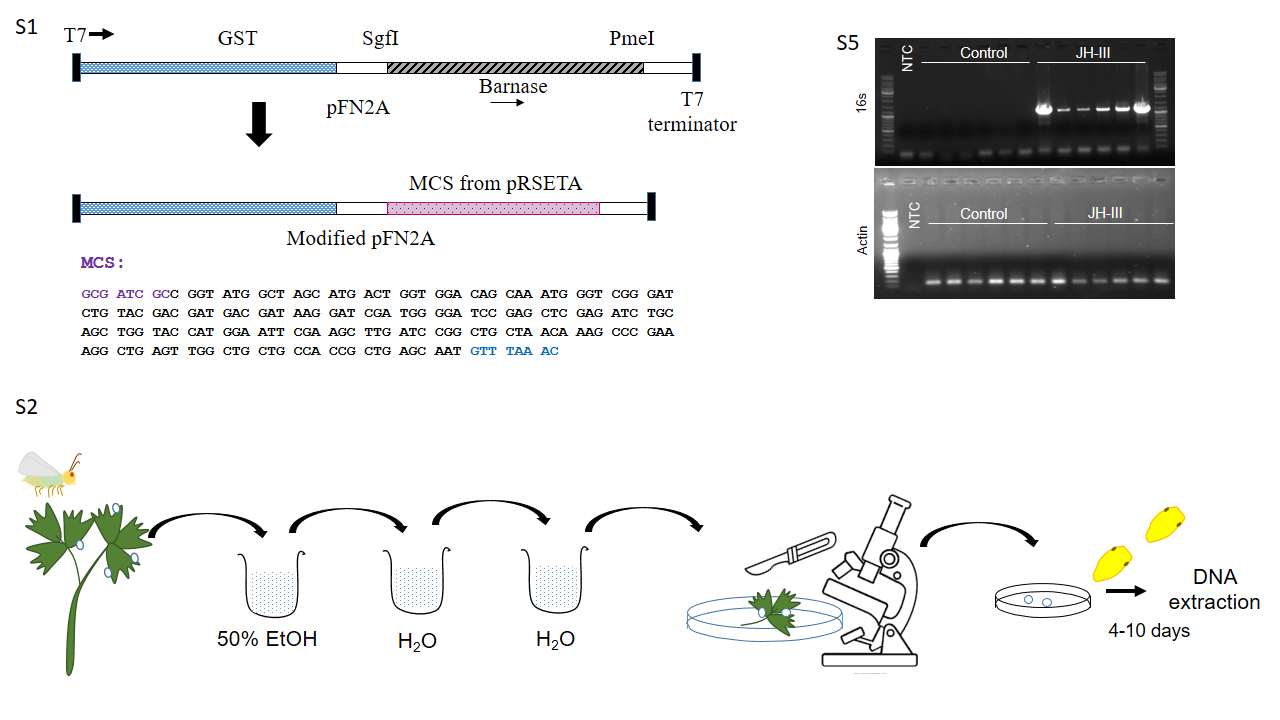

**Fig. S1.** Procedure by which eggs were separated from the leaves and were allowed to hatch in a sterile environment for CLso detection.

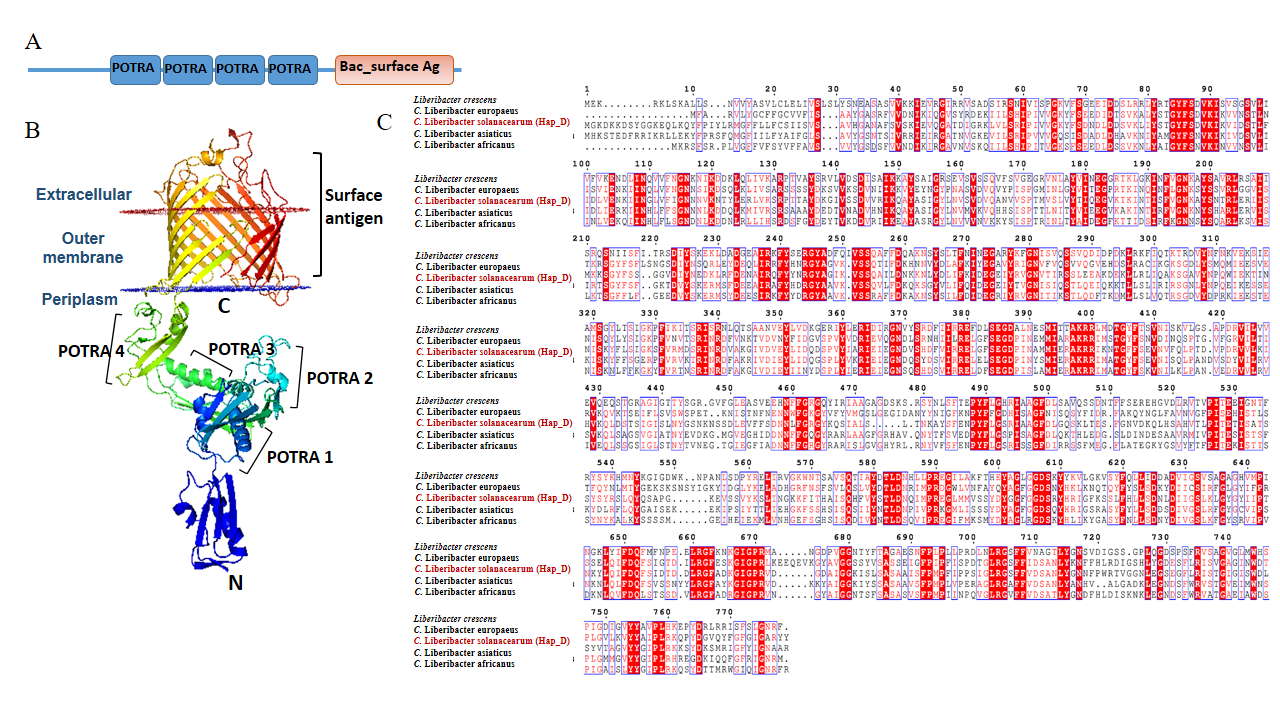
**Fig. S2.** Domain and conserved sequence analysis of OmpA. **A** and **B**, Presence of four polypeptide transport-associated domain (POTRA) and an extracellular surface antigen domain is conserved in Liberibacter OmpA. **C**, Multiple sequence alignment of known Liberibacter OmpA proteins showing conserved residues in red.

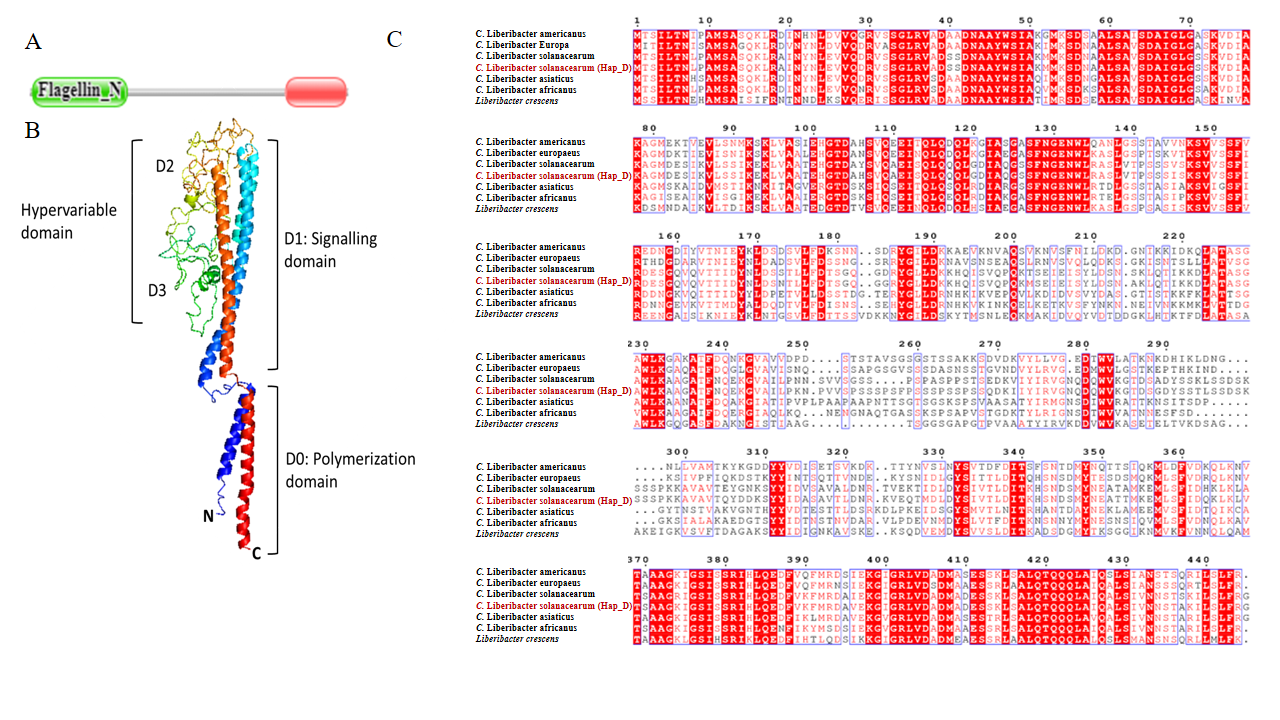

**Fig. S3.** Domain analysis of Flagellin. **A** and **B**, Flagellin structure showing signaling and polymerization domains. **C**, sequence alignment showing conserved residues across all known Liberibacter flagellins.

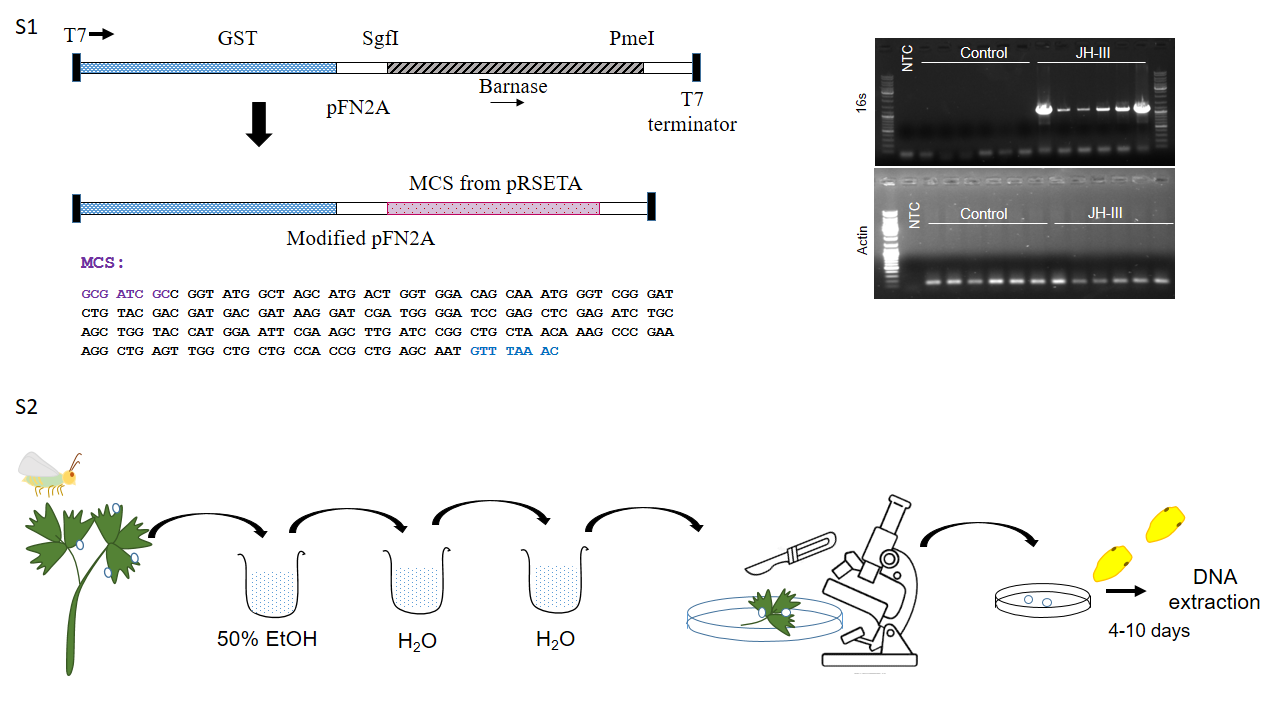

**Fig. S4.** PCR detection of CLso using 16s-rRNA specific primers and Actin as housekeeping control showing absence of CLso in L+ control ovaries and presence of CLso in JH-III treated ovaries.
